# The ideotype for drought tolerance in bioenergy *Populus nigra*

**DOI:** 10.1101/2024.02.07.579233

**Authors:** Hazel K Smith, Jaime Puértolas, Cyril Douthe, Giovanni Emiliani, Alessio Giovannelli, Libby S Rowland, Mike Allwright, Jack H Bailey-Bale, Pili M Valdes-Fragoso, Elisabeth K Larsen, Giorgio Alberti, Alessandro Zaldei, Andrew D Hirons, Franco Alasia, Miquel Ribas-Carbo, Jaume Flexas, Ian C Dodd, William J Davies, Gail Taylor

## Abstract

Fast-growing perennial trees such as Populus nigra L. are important species for wood, plywood, pulp, and bioenergy feedstock production, yet tree vigor in a changing climate is poorly understood. This research aimed to identify breeding targets for yield in water-limited environments, alongside unraveling the relationship between drought, yield, and glucose release in P. nigra. A diversity panel of 20 P. nigra genotypes, selected from a wide natural association population, was grown at three divergent European sites. Through extensive phenotyping of physiological and morphological productivity and water-use traits, under irrigated conditions and when exposed to a progressive drought, we elucidated the adaptive and plastic drivers underlying tree productivity. We have identified the underpinning traits for drought tolerance, whereby high yields can be maintained under water deficit, in this key species. This highlighted the importance of examining the yield stress index (YSI) over the drought resistance index (DRI) to assess genotypes for performance under moderate drought. In this way, we found genotypes with high hydraulic capacity, and large leaves made up of many cells to be best suited to multiple European environments, with contrasting water availability. Moreover, we identified genotypes that combine yield and water use efficiency, with good glucose release potential, which will be important traits for the future of poplar as a bioenergy crop. Vigorous poplar genotypes, which are adapted to wet climates showed high environmental plasticity. However, in European drought scenarios, these trees outperform drought resistant genotypes, and some exhibit good glucose release. These trees are a valuable resource for the future.

## Introduction

Alongside rising temperatures, it is predicted that the frequency and severity of droughts will increase across Europe in the coming years (Lindner *et al*. 2010; IPCC, 2021, and in temperate regions, these drought episodes have been linked to long-term reductions in tree growth and increased forest die-back (Breda *et al*., 2006, Anderegg *et al*., 2013, Greenwood *et al*., 2017). In addition to the impacts of climate change, human population growth will increase food, fuel, and fibre demands (Spiertz *et al*., 2009). *Populus* shows some of the highest growth rates for temperate trees (Taylor, 2002) and is widely grown across the northern hemisphere for wood, plywood, and pulp, and as a dedicated bioenergy crop (Zsuffa *et al*., 1996; Allwright and Taylor, 2016). Although the genus can be grown on marginal land, thus reducing competition with food production, high productivity requires significant water use (Barigah *et al*., 1994). However, *Populus* spp. are diverse with wide genetic variability, and trees of this genus show markedly different responses to water deficits (Bonhomme *et al*., 2009; Monclus *et al*., 2005, 2009; Bogeat-Triboulot *et al*., 2007; Viger *et al*., 2016). Capturing this variation is important so as to improve climate resilience across this important genus.

Predicted drier climates necessitate the development of *Populus* genotypes, which are able to tolerate moderate water deficits without suppression of carbon assimilation or subsequent productivity (Monclus, 2006; Hamanishi *et al*., 2015). If these trees are to be used for bioenergy, saccharification potential, and glucose release must also be maintained or enhanced (Brethauer and Studer, 2015; Karimi and Taherzadeh, 2016). Previous studies with hybrids have identified genotypes that combine high productivity, WUE (the amount of biomass gained per unit of evapotranspiration), and drought tolerance (Marron *et al*., 2005; Monclus *et al*., 2005, 2006, 2009), yet none of these have considered saccharification potential and all examined non-European germplasm. Here, we use a genetically diverse panel of *Populus nigra* L., selected from across contrasting European riparian zones (Rohde *et al*. 2011; Viger *et al*., 2016), to elucidate the intricate basis of a wide-range of adaptive drought responses and their level of plasticity, as well as their relationship with yield and saccharification potential.

To date, robust breeding targets for drought tolerance have been elusive as such traits vary, depending on the timing, duration and severity of the water deficit (Bréda *et al*., 2006; Tardieu & Tuberosa, 2010, Tardieu et al, 2011). Plants mitigate against limited soil water availability through a range of mechanisms and breeding for drought tolerance should be specific to the targeted crop system and the type of drought that occurs most frequently (Tardieu, 2011). This implies targeting the traits which are most relevant in maintaining productivity in that crop system and, for *Populus*, this means maximising overall biomass accumulation. The drought scenario, which is predicted for *Populus* in Europe, combines moderate water deficits with a requirement for sustained productivity. Thus, drought tolerant poplar genotypes must maintain cell production, expansion, leaf growth, and gas exchange rates under soil water deficits (traits identified as being drought responsive in Liu & Dickmann, 1992; Tschaplinski *et al*., 1998; Marron *et al*., 2002; Marron *et al*., 2003; Tardieu & Tuberosa, 2010; Brunner *et al*., 2015; de Ollas & Dodd, 2016), while other traits/processes important during very intense drought events like cell protection, lignification, reduced vessels lumen area (related to cavitation prevention) (Gebre *et al*., 1994; Marron *et al*., 2002; Sack and Holbrook, 2006; Cochard *et al*., 2007, Jacobsen et al, 2019) may be less important.

In general, two main drought response strategies have been identified in *Populus* – tolerance and avoidance (Marron *et al*., 2003; Monclus *et al*., 2006; Giovannelli *et al*., 2007). The former relies on maintaining stomatal conductance and photosynthesis, while limiting tissue dehydration through osmotic adjustment (Gebre *et al*., 1994; Marron *et al*., 2002; Barchet *et al*., 2013; Traversari *et al*., 2018). The latter minimizes water loss through rapid stomatal closure, leaf growth inhibition, and/or leaf abscission (Couso and Fernandez, 2012). The combination of traits that underlie drought responses are often adaptive and develop as a consequence of environmental conditions of the native origin of a genotype, with populations exhibiting wide natural variation (Viger *et al*., 2016). However, alongside adaptive drought responses, plants often exhibit plasticity in their responses. Phenotypic plasticity occurs in response to changing environmental conditions (Bradshaw, 1965) and is of critical importance to plants due to their sessile nature. The extent of plasticity in response to the environment is variable; dependent on both genotype and trait, and high plasticity could enable populations or individuals to survive, or maintain performance, in a rapidly changing climate (Benito Garzόn *et al*., 2010). To make genetic gains in *Populus*, these target traits and their plasticity must be thoroughly understood. Thus, we have undertaken a detailed morphological and physiological screening of a diverse panel of genotypes both across sites, and in response to varying water availability.

In these long-lived species, productivity can only be assessed at the end of the rotation age. It is therefore important to identify traits that rapidly and cost-effectively predict drought tolerance and yield. Stem volume and individual leaf area correlate positively with yield and are robust early indicators of productivity (Rae *et al*., 2004; Marron and Ceulemans, 2006; Monclus *et al*., 2005, 2006). Alongside this, it has been reported that trees with lower specific leaf area show higher drought resistance and lower mortality (Greenwood *et al*., 2017). However, other leaf traits such as cell patterning are less reliably linked to final biomass due to high environmental regulation (Allwright and Taylor, 2016). WUE itself can be estimated through carbon isotope discrimination (Δ^13^C) of plant tissues (Farquhar & Richards, 1984; Farquhar *et al*., 1989; Bonhomme *et al*., 2008) and is an attractive breeding target given it is a highly heritable, integrative measurement (Marron et al, 2014). To increase water productivity (ratio of yield to water use), WUE must be increased without sacrificing biomass production. However, the relationship between WUE and yield is variable (Monclus *et al*., 2005, 2006; Prasolova *et al*., 2003; Rae *et al*., 2004), making it an uncertain general breeding target at present. It is therefore important that we assess the link between WUE and yield in *P. nigra* as well as considering other traits to identify novel high-throughput targets.

This study aims to identify breeding targets for yield in water-limited conditions, while also characterizing the relationship between drought, yield, and saccharification potential in *P. nigra*. Thus, we evaluated the relative importance of genetic adaptation and phenotypic plasticity of growth traits under water deficit. To this end, a diversity panel of 20 *P. nigra* genotypes, selected from a wide natural association population, was examined at multiple European sites under irrigated conditions and exposed to a progressive drought at one northern Italian site, to assess productivity, bioenergy feedstock quality and phenotypic responses.

## Materials and methods

### Plant material and experimental design

A diversity panel of twenty *P. nigra* genotypes were selected from a natural, wide population of 661 genotypes (described previously by Rohde *et al*., 2011; Allwright *et al*., 2016; Faivre-Rapant *et al*., 2016; Viger *et al*., 2016). The population consisted of genotypes originating from Spain, France, Italy, Germany and The Netherlands (Figure 1A) and the diversity panel represented the full range of WUE within each country of origin, estimated from wood Δ^13^C (Δ^13^C_wood_) (Farquhar and Richards, 1984) determined in a previous Belgian common garden experiment (Viger *et al*., 2016, Figure 1B–C). The diversity panel became 16 genotypes, as four were excluded due to poor establishment in Italy, while the rest of the panel was subjected to extensive phenotyping (as described in Taylor *et al.,* 2019), to determine the physiological basis of drought tolerance in black poplar. From this point forth, the diversity panel will refer to the 16 genotypes that were tested in all three locations.

**Figure 1.**
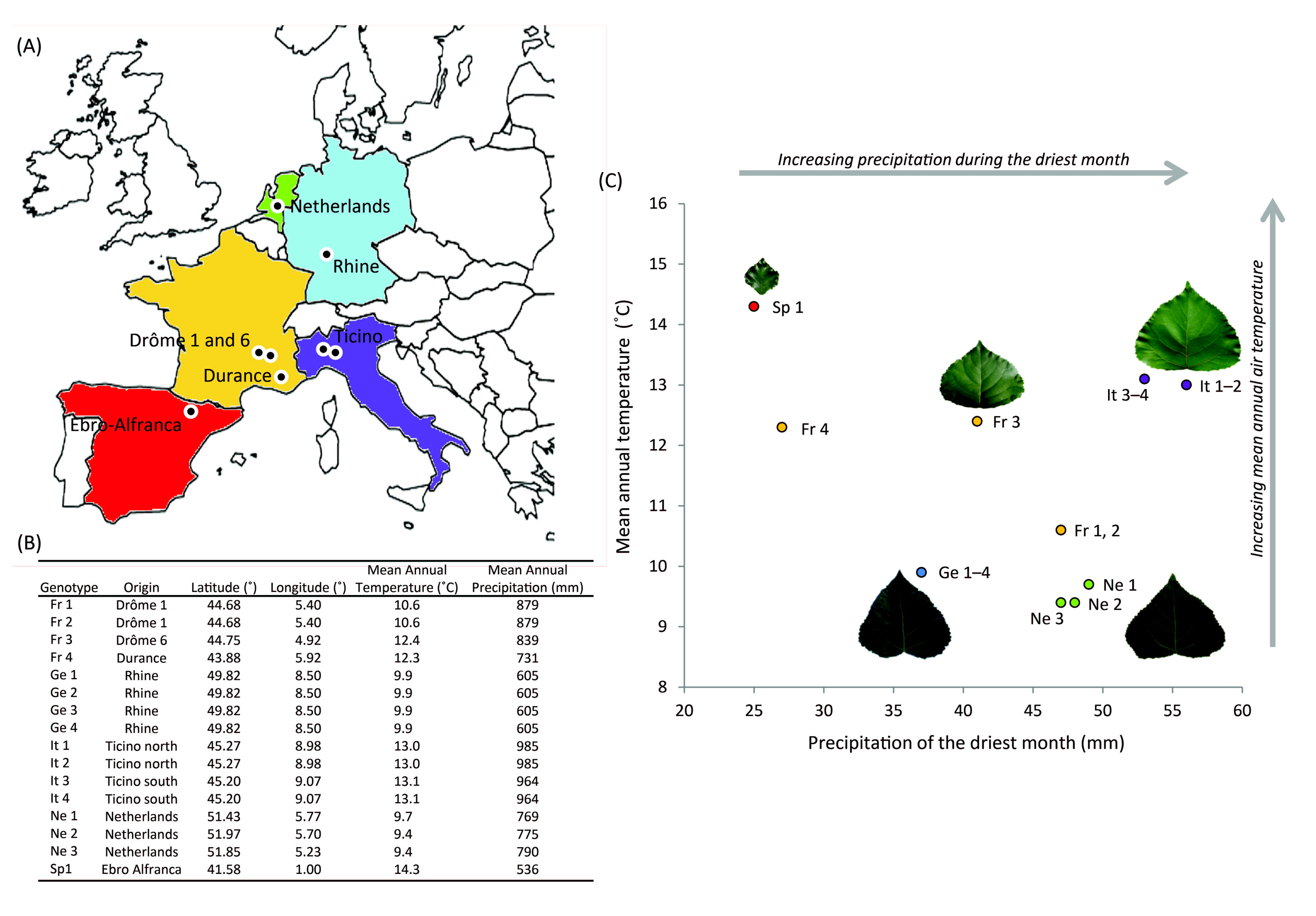
Description of the diversity panel. Genotypes were selected from a wide European association population of *Populus nigra* L, originating from distinct populations (A), with varying environmental conditions (B–C).

The wide association population was planted from hardwood cuttings at a field site in northern Italy (44.6 °N, 7.6 °E, 311.8 m asl) in Apr. 2013, following a randomised block design of eight blocks, each of which was surrounded by two rows of a commercial *Populus* hybrid ‘Monviso’ to counteract edge effects. Each block contained one replicate of each genotype and these were randomly positioned at a spacing of 0.75 x 2.3 m. Blocks were spaced 5 m apart in a 2 x 4 arrangement. All blocks received commercial drip irrigation, weed control and insecticide treatments from Apr. 2013–Dec. 2015. Each winter, trees were coppiced at the base before being single-stemmed at the start of each growing season, following commercial practice. The site was instrumented with a meteorological station, which monitored precipitation, temperature and relative humidity. Two Decagon EC5 soil moisture probes (Decagon Devices, Washington, USA) were installed in the center of each block at 20 cm and 40 cm depth in mid-Jun 2015, with remote data transmission and logging.

### Wide association population trials

Tree growth (stem volume index, SVI) and individual leaf area of each genotype in the diversity panel were compared between the previously described trial in Italy and two further sites in Europe: Belgium, Geraardsbergen (50.8 °N, 3.9 °E, 26.1 m asl) (Viger *et al*., 2016) and the UK, Hampshire (51.1 °N, -1.2 °E, 95.7 m asl) (Allwright *et al*., 2016). This presented a novel opportunity to assess plant performance and drought response across a range of European environments. Furthermore, Δ^13^C_wood_ discrimination was also compared between the Belgian and Italian trials.

### Irrigation regime at the Italian field site

The soil type at the Italian field site was loamy with a porosity of 40% and a density of 1.58 g cm^-3^. The meteorological daily means were: 18.75 °C temperature; 24.07% relative humidity and 0.14 mm precipitation. Following pre-season measurements (completed on 7 Jun. 2015), irrigation was withheld in blocks 1–4 (drought), while blocks 5–8 were irrigated using a drip irrigation system. The effect of the drought treatment on soil moisture was determined using the B.A.C.I. (Before-After-Control-Impact) design (Underwood, 1993). Three weeks were allowed for probe stabilisation (from installation of the probes in mid-Jun. until 4 Jul. 2015). The period of 4–19 Jul. was considered as the “before” period, and from 20 Jul.–4 Aug. – corresponding to the period of physiological measurements – as the “after” period for the B.A.C.I. design. For each of the treatments, the soil moisture (SM) response is then expressed as:

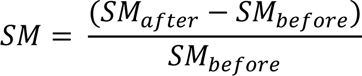

### Biomass measurements

Immediately before applying the irrigation regimes, allometric biomass measurements were taken for the diversity panel. These measurements were taken over three consecutive days during the first week of Jun. 2015 and consisted of main stem height and diameter at 22 cm from the ground. SVI was calculated as:

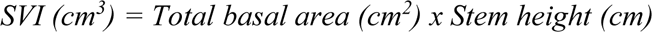

After eight weeks of the irrigation regime, allometric measurements were repeated. Additionally, the first mature leaf from each tree of the diversity panel was harvested in the last week of Jul. 2015. This leaf was imaged on a scaled background using a digital SLR camera before being analysed in ImageJ software (http://rsb.info.nih.gov/ij/) (Abramoff *et al*., 2004) for individual leaf area, length and width. These leaves were then oven-dried at 80 °C until constant weight was achieved.

### Leaf cell patterning

Epidermal imprints were taken from the abaxial surface of the first mature leaf of each tree as previously described by Gardner *et al*. (1995) in the last week of Jul. 2015. Epidermal imprints were stored at 3 °C, before imaging at 20x magnification on an Axioplan Zeiss microscope (Zeiss, Oberkochen, Germany). The images were assessed in Image J software and the cell area of 10 randomly selected cells per imprint were measured. A 5000 µm^2^ grid was applied to the image and epidermal cells were counted in five randomly selected grid squares. The number of stomata in the full image was counted. Stomatal density (stomatal number per unit area), stomatal index (SI, ratio of the number of stomata per total cell number as a percentage), cell density and cell number per leaf (CNPL, cell density x individual leaf area) were calculated.

### Gas exchange measurements

Gas exchange measurements were performed using a Li-6400 equipped with a 6 cm^2^ open-top chamber on 2 and 3 Aug. 2015 (four blocks per day). The youngest fully-expanded leaf on the main stem was measured for each replicate of the diversity panel genotypes. CO_2_ concentration surrounding the leaf (C_a_) was controlled at 400 µmol mol^-1^ via the Li-6400 CO_2_ injector. Leaf temperature and water vapour concentrations were not controlled so as to measure leaf performance in field conditions. All measurements were made at 1430–1630 h on leaves which were fully exposed to the sun before and during the measurement (PPFD>1500 µmol photons m^-2^ s^-1^) (Zhang and Gao, 2000). Each leaf was clamped, while assimilation and stomatal conductance stabilised (∼2 min), before data acquisition with five readings at 2–3 s intervals. These readings were averaged to obtain leaf photosynthetic parameters. At the same time, an Optris LS LT portable laser thermometer (Optris Infrared Thermometers, Berlin, Germany) was used to make a point measurement of leaf temperature of a sunlit, mature leaf for each tree.

### Leaf water potential at predawn and midday

Predawn leaf water potential (Ψ_pd_) was measured on the two nights previous to gas exchange (1 and 2 Aug. 2015), and four blocks were measured each night. The afternoon previous to the measurements, a mature leaf from each tree was wrapped in aluminum foil to prevent night transpiration, allowing the leaf water potential to closer equilibrate with soil water potential and likely reflect the water potential in the stem, where the petiole is attached. Predawn measurements were made at 0300–0430 h while midday water potential (Ψ_md_) was measured at the same time as gas exchange, on a sunlit mature leaf located in the upper third of the main stem. Immediately after excision, each leaf was placed inside a dark plastic bag to minimise water loss and time between excision and measurement was <5 min.

### Osmotic potential at full turgor

Samples were collected in the last week of Jul. 2015 alongside biomass and physiological measurements. A mature branch was excised from each tree and the cut surface was placed into water. Subsequently (2–3 min), the stem was submerged and re-cut at least two internodes towards the apex before being re-hydrated to full turgor overnight in a humid, dark box at room temperature. The following day, a leaf disc (7 mm diameter) was taken from a fully expanded leaf. Discs were wrapped in aluminum foil and snap frozen in liquid nitrogen before being stored at -80 °C.

Leaf solute concentration was measured as in Sjöman *et al*. (2015) using a vapour pressure osmometer (Vapro 5600, Wescor, Logan, USA). Solute concentration (mmol kg^−1^) was measured after 10 min equilibration. Osmotic potential was calculated according to the equation:

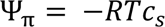

Where R is the gas constant, T is temperature in Kelvin, and c_s_ is the solute concentration in mol m^3^ (=mmol kg^-1^).

### Carbon isotope analysis

A mature, sunlit leaf from each tree was collected the day before gas exchange measurements (1 Aug. 2015, 1430–1730 h). Leaves were excised from the main stem, just below the leaf used for gas exchange measurements, and placed in a 2 ml eppendorf tube. The tubes were immediately frozen in liquid nitrogen and stored at -20 °C before being freeze-dried and finely ground. Wood samples were collected in Dec. 2015 from the upper quarter of the main stem. Debarked wood samples had their pith removed before being oven-dried to constant mass at 105 °C. These samples were then ground to a fine powder using a mixer mill (MM300, Retsch, Haan, Germany). Samples of both leaves and wood were collected from material which was produced during the time when the drought treatment was imposed. For leaf and wood samples, 2 mg was transferred into tin capsules, per tree. Samples were then combusted in an elemental analyser (Thermo Flash EA 1112 Series, Bremen, Germany), and directly injected into a continuous-flow Isotope Ratio Mass Spectrometer (Thermo-Finnigan Delta XP, Bremen, Germany) for isotope analysis. Peach leaf standards (NIST 1547) were run every six samples. The standard deviation of the analysis was always below 0.1‰. Carbon isotope composition (δ^13^C) was then calculated relative to the Pee Dee Belemnite standard as in Farquhar *et al*. (1989).

### Leaf abscisic acid (ABA)

The same ground leaf material prepared for carbon isotope analysis had deionised water added to it at a 1:70 weight ratio. The aqueous extract was incubated overnight at 4 °C in a shaker. Leaf ABA concentration was measured using a radioimmunoassay (Quarrie *et al*., 1988). A dilution spike recovery test (Zhang and Davies, 1990) showed no immunoreactive contamination for black poplar leaf tissue (data not shown).

### Leaf ethylene

Samples for ethylene evolution determination were collected on 2 and 3 Aug. 2015 (1130**–**1200 h, 1730**–**1800 h). Leaves (0.5**–**1.5 g in fresh weight) were sampled adjacent to those used for gas exchange, Ψ_md_ and ABA analysis. Leaves were placed in 24 ml glass tubes with an airtight rubber seal and allowed to incubate for 60 min in darkness and then 45 min under a 60 W lamp. After the incubation period, 2 ml of air was extracted from the tubes with a syringe and injected into a 20 ml vial through a septum screw cap. Vials were transported to the lab and, two days after extraction, total ethylene in the vial was measured with an ethylene detector (ETD-300, SensorSense, The Netherlands). Leaves were oven-dried at 70 °C for 48 h and fresh weight was estimated using an equation calculated for black poplar leaves in a greenhouse experiment (data not shown). Ethylene evolution (nl h^-1^ g^-1^) was calculated assuming that the 2 ml of air was representative of the 24 ml vial as follows:

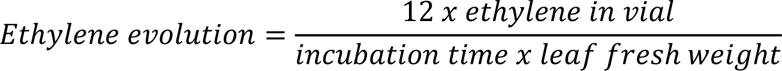

### Saccharification Potential

The 150–850 µm fraction of the ground wood material used for Δ^13^C was assayed for saccharification potential (as described in Allwright *et al*., 2016). Briefly, moisture content was calculated from weight loss of an aliquot of each sample after oven-drying to constant weight at 105 °C. A 10 mg sample of un-dried powder underwent acid pre-treatment and ethanol wash steps followed by 48 h saccharification with fungal cellulose and cellobiase enzymes (Sigma, Missouri, USA) at 55 °C in a rotating thermomixer. The supernatant was assayed with GOD-POD solution (Van Acker *et al*., 2013) and saccharification potential was calculated as the sample glucose yield as a percentage of post pre-treatment oven-dry weight.

### Wood basic density, xylem morphology and theoretical hydraulic conductivity calculation

In Dec. 2015, woody stem discs of 8**–**10 cm in thickness were collected from the main stems at 80 cm height, kept in plastic bags and transported to the laboratory at 4 **°**C.

#### Wood basic density

For each disc, two prismatic sub-samples were taken from the central part of the sample (excluding pith and bark) following the radial, tangential and longitudinal directions (20 x 20 x 20 mm). Each sub-sample was composed of the current annual ring. The samples were oven dried at 103 °C for 96 h and dry weight (DW, kg) measured (BP110S, Sartorius, Goettingen, Germany; 0.1 mg accuracy). The dry volume (DV, m^3^) was measured by water displacement (Berta *et al*., 2010) and basic density (BD, kg m^-3^) calculated as reported by Carriero *et al*. (2015).

#### Xylem morphology

Two to three cm sections of stem were cut from the woody disc for each tree. These portions of the stem were then placed in a 50:50 mixture of ethanol and water and stored at 5 °C. The stem sections were then fixed through ice on a Peltier plate, and transverse sections of 8–12 μm thickness were cut using a rotary microtome. The sections were stained with a solution of 0.04% safranin, 0.15% astrablue and 2% acetic acid in distilled water (Emiliani *et al*., 2011), and permanently fixed with a histological mounting medium (Eukitt, Bioptica, Milan, Italy). A Nikon Eclipse 800E light microscope connected to a Nikon DS-Fi2 microscope camera (Nikon Corporation, Tokyo, Japan) was used for anatomical observations. Digital images of the cross-sections were then analysed and transversal stem structure examination was performed on 4–6 independent images per sector using the Nikon Nis-Elements software. For each image, the vessel density (N_v_, mm^-2^) and mean vessel lumen cross-section (A_v_, μm^2^) were measured. The vessel and fibre element length (L_VE_ and L_FB_ respectively, μm) were determined after maceration of wood slivers using a Jeffrey’s solution for 8 h at room temperature. After two washing steps, cell separation took place in a droplet of water. Cell length was measured for at least 50 vessels and 100 fibres per woody ring as proposed by Scholz *et al*. (2013).

#### Theoretical Hydraulic Conductivity (K_st_)

The diameter of each vessel was calculated as the diameter of a circle with an area equivalent to the mean lumen cross-section area. The frequency distribution of xylem vessels was then categorised by 10 μm diameter classes. The hydraulic weighted vessel diameter D_h_ was calculated following Sperry and Saliendra (1994) as:

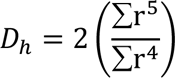

Where r is the radius in μm. Calculation of D_h_ incorporates the disproportionate contribution of large vessels to total flow and gives the average diameter needed for a given vessel density to result in the theoretical hydraulic conductivity for that stem (Tyree *et al*. 2002).

K_st_ was calculated following Santiago *et al*. (2004) using the Hagen-Poiseuille equation for ideal capillaries assuming laminar flow:

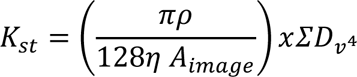

where ρ is the density of water (998.2 Kg m^-3^ at 20 °C); η is the viscosity of water (1.002 x 10^-9^ MPa s at 20 °C); A_image_ is the area of the image analysed (m^2^) and D_v_ is the mean vessel diameter (μm).

### Statistical analysis

Correlations were conducted in Minitab (Minitab 17.3.1, Pennsylvania, USA) and were based on linear models with the Pearson’s coefficient of correlation (r) given. Principle Component Analysis (PCA) was conducted in R (R Core Team, 2015) using the package FactoMineR and the function PCA, and PCA plots were made using ggplot2. All relationships and mean comparisons were accepted as significant at p value<0.05.

The Drought Resistance Index (DRI, Fischer and Maurer, 1978) and Yield Stress Index (YSI, Fernandez, 1992) of each genotype and trait combination were calculated using the following equations:

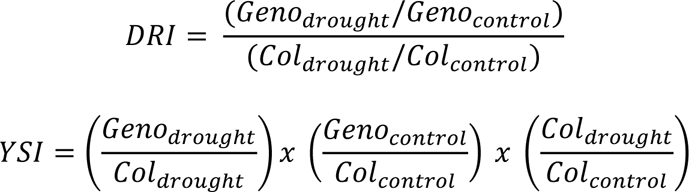

Where *Geno* indicates the genotype mean trait value, and *Col* denotes the collection mean trait value.

Drought resistance index is a calculation of the reduction in yield (biomass) when a genotype is under water stress, compared to the mean reduction across the rest of the population being studied. The yield stress index is able to identify genotypes that display high yields under both water stressed and control conditions. It can therefore indicate genotypes with the greatest drought tolerance and yield potential, which can otherwise be difficult to pinpoint when considering DRI alone.

## Results

Soil water content decreased during the period of drought application, with negative B.A.C.I. values, and this reduction was approximately 15% in the drought treatment, while it was less than 5% for the irrigated plots (p<0.05, Figure S1). To the best of our knowledge, all genotypes had access to water at the same soil depths.

### Phenotypic plasticity across three European common garden sites

Tree productivity showed significant variation across the diversity panel with Sp1 and Fr1 genotypes showing the lowest growth at the Savigliano field site, Italy (Figure 2A). Although the variability of both individual leaf area and tree height was site-dependent, the strong correlation between the two was maintained across sites (Figure 2B). Further, the genotypes with high individual leaf areas, which were the most productive trees, generally had the highest phenotypic plasticity for leaf area (Figure 2C). Values of Δ^13^C_wood_ were lower and genotypic variability higher when trees were grown at the Italian site when compared to the Belgian common garden. Δ^13^C_wood_ was negatively and significantly correlated with individual leaf area in Italy (r=-0.72, p<0.002), but this trend was not significant in Belgium (Figure 2D). Despite the phenotypic plasticity observed between sites, the ranking of Δ^13^C_wood_ values were largely conserved between the two sites (r=0.62, p<0.05, Figure 2E). As with individual leaf area, the genotypes with the highest values of Δ^13^C_wood_ in Belgium generally experienced the greatest decrease when grown in Italy (Figure 2D). Mean individual leaf area increased with decreasing experimental latitude, yet this was not significantly related to precipitation during the driest month at the native site (Figure 2F).

**Figure 2.**
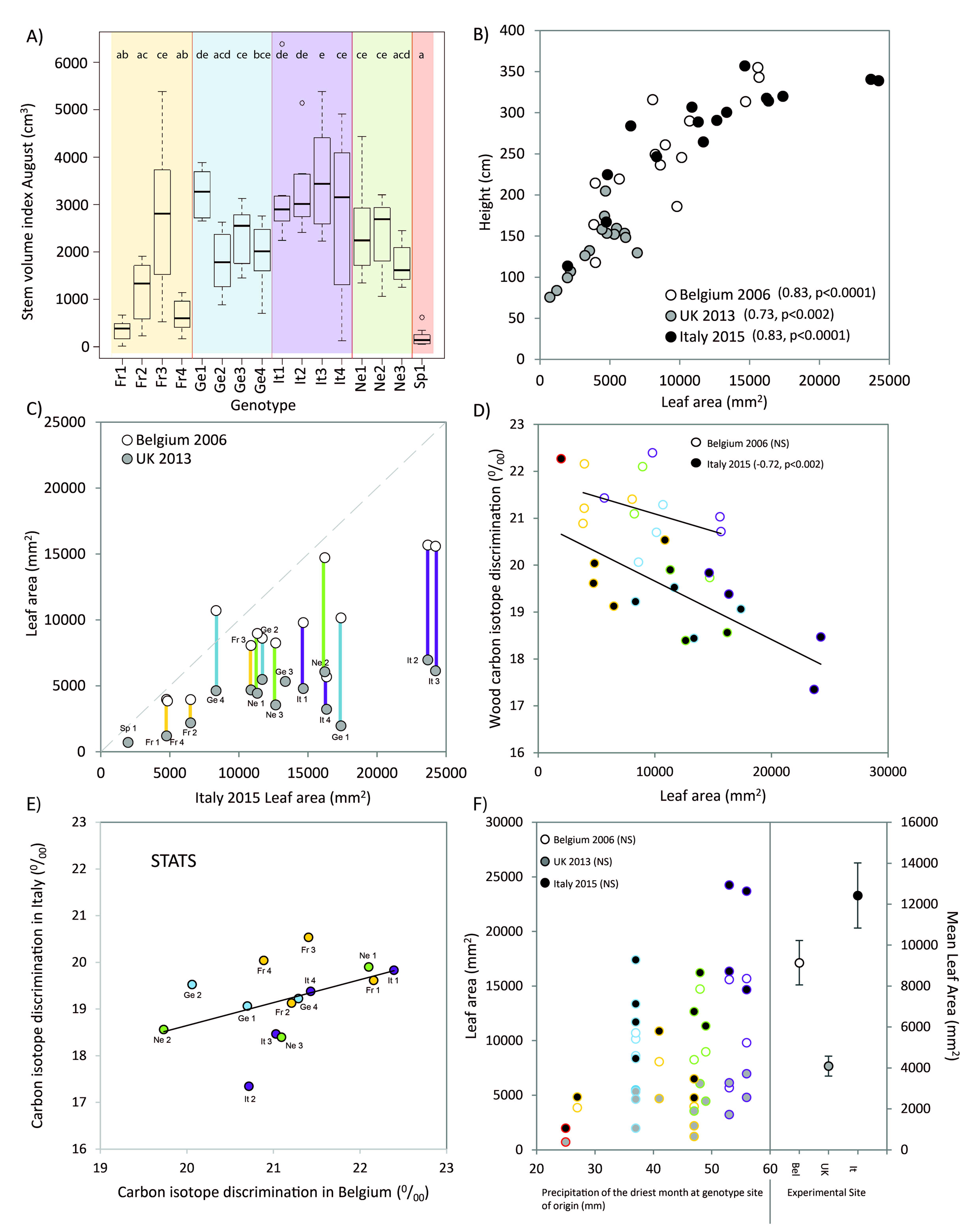
Performance of the diversity panel grown at contrasting European sites. Stem volume index (SVI) in Savigliano, Italy for each of the selected diversity panel genotypes (A); individual leaf area versus height at the three European experimental sites, Belgium, Italy and the UK, (B); individual leaf area in Belgium and the UK versus individual leaf area in Savigliano, Italy (C); wood carbon isotope discrimination versus individual leaf area in Belgium and Italy (D); wood carbon isotope discrimination in Belgium and Italy (E); precipitation of the driest month at the site of origin versus individual leaf area in Belgium, Italy and the UK, as well as, mean leaf area for the three European experimental sites (F).

### The physiological and morphological basis of tree growth

To identify the most important correlations underlying tree growth and drought responsiveness, correlations between traits were examined (Figure 3, Table S1). When the performance of both control and droughted trees were combined (Figure 3A), yield (SVI) was positively linked to leaf morphology with large trees producing large leaves comprising many cells (individual leaf area, r=0.88, p<0.0001; CNPL, r=0.86, p<0.0001; stomatal index, r=0.43, p<0.05). These large trees also displayed high hydraulic capacity (D_h_, r=0.64, p<0.0001; K_st_, r=0.66, p<0.0001; vessel diameter, r=0.80, p<0.0001). However, high productivity was linked to reduced Δ^13^C_wood_ (r=-0.56, p<0.001) and saccharification potential (r=-0.47, p<0.01).

**Figure 3.**
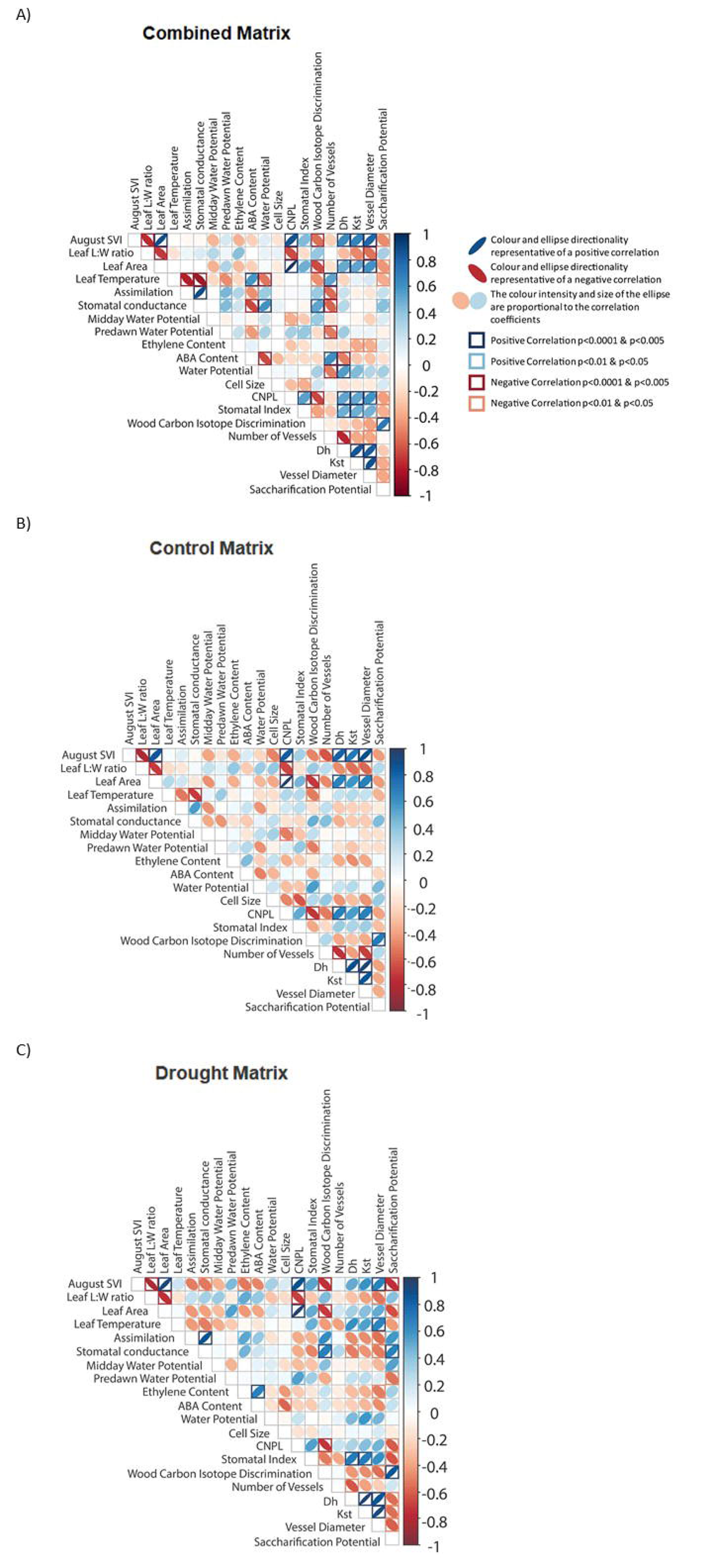
Correlation matrices for a subset of physiological and morphological traits when the diversity panel was grown in Italy. Irrigated and droughted regimes pooled (A), control (B), and drought (C). Correlations for all traits can be seen in Table S1.

Leaf and xylem hydraulic traits were generally correlated to productivity (SVI) (individual leaf area, r=0.84, p<0.0001 and r=0.93, p<0.0001; CNPL, r=0.84, p<0.0001 and r=0.88, p<0.0001; stomatal index, NS and r=0.52, p<0.05; D_h_, r=0.82, p<0.0001 and NS; K_st_, r=0.70, p<0.005 and r=0.59, p<0.05; vessel diameter, r=0.89, p<0.0001 and r=0.68, p<0.005) under both control (Figure 3B) and drought (Figure 3C) irrigation regimes. However, SVI was negatively related to Δ^13^C_wood_ (r=-0.74, p<0.001) and saccharification potential (r=-0.74, p<0.001) only under drought conditions. Further, although saccharification potential showed only one significant relationship under the irrigated regime (to Δ^13^C_wood_, r=0.69, p<0.005), the imposition of drought elucidated a number of other associations. In drought conditions, trees with high saccharification potential tended to be lower yielding (r=-0.74, p<0.001), made up of smaller leaves, with a lower cell number per leaf and stomatal index (individual leaf area, r=-0.65, p<0.01; CNPL, r=-0.62, p<0.01; stomatal index, r=-0.63, p<0.01). These high saccharification potential trees also demonstrated a reduced hydraulic capacity (D_h_, r=-0.55, p<0.05; K_st_, r=-0.56, p<0.05; vessel diameter, r=-0.60, p<0.05). Drought also highlighted that these high saccharification potential trees displayed higher values of Δ^13^C_wood_ (r=0.86, p<0.0001).

Unlike saccharification potential, Δ^13^C_wood_ was more consistently linked to tree morphology irrespective of the irrigation regime. Under both conditions, Δ^13^C_wood_ was negatively linked to individual leaf area (r=-0.70, p<0.005 and r=-0.72, p<0.005) and CNPL (r=-0.73, p<0.001 and r=-0.72, p<0.005). The positive relationship between Δ^13^C_wood_ and saccharification potential was relatively stable between conditions (r=0.69, p<0.005 and r=0.86, p<0.0001). However, the physiological links were less stable with CO_2_ assimilation and stomatal conductance tending to be higher in trees with high Δ^13^C_wood_ under the irrigated regime (r=0.63, p<0.01 and r=0.68, p<0.005 respectively), but unrelated under drought. On the other hand, predawn water potential was negatively correlated with Δ^13^C_wood_ under drought (r=-0.50, p<0.05) but the two were not linked in the irrigated regime. Conversely, trees with high osmotic potential at full turgor tended to also have high Δ^13^C_wood_ values under drought (r=0.55, p<0.05), but not in control.

While the native latitude did not relate to SVI, longitude and altitude of the genotype’s origin were related to productivity. In both control and drought, the more productive trees tended to originate from more easterly longitudes (r=0.70, p<0.005 and r=0.66, p<0.005) and lower altitudes (r=-0.68, p<0.005 and r=-0.75, p<0.001). Trees from lower latitudes and longitudes, such as the Spanish genotype, exhibited higher Δ^13^C_wood_ (r=-0.67, p<0.005 and r=-0.64, p<0.01 respectively) and saccharification potential (r=-0.62, p<0.01 and r=-0.64, p<0.01 respectively). High Δ^13^C_wood_ and saccharification potential was linked to native sites of higher altitude (r=0.81, p<0.0001 and r=0.79, p<0.0001 respectively).

PCA was employed to identify the drivers of these trait interactions (Figure 4). The same subset of variables was used for this analysis as for Figure 3, with highly correlated traits removed. The first axis of the PCA explained 27.92% of the variance, and the second 17.24%. The first axis (F1) was strongly related to biomass traits and region of origin (Figure 4A). Thus, F1 tended to separate genotypes, with productivity increasing from left to right of the axis. Conversely, F2 was associated with physiological variables and separated the irrigation regimes, with droughted plants on the upper part and irrigated trees on the lower part of the axis (Figure 4B). Interestingly, both Δ^13^C_wood_ and saccharification potential fell between the main planes. F1 showed high values of individual leaf area, and cell number per leaf were associated with more productive genotypes (high SVI), and higher theoretical hydraulic conductivity (K_st_). Increased SI, latitude and longitude of the population of origin were also associated with more productive genotypes, but to a lesser extent. Italian genotypes (located on the right side of the plane, particularly It2 and It3) were associated with the highest values of biomass production, while the Spanish and French genotypes showed lower productivity. Genotypes from the Netherlands and Germany were closely grouped together, on the high productivity side of the F1 main plane. The second main plane of the PCA (F2), which was related to physiological variables (Figure 4B) and correlated most strongly with variables such as gas exchange (A, stomatal conductance and leaf temperature), and wood anatomy (number of vessels).

**Figure 4.**
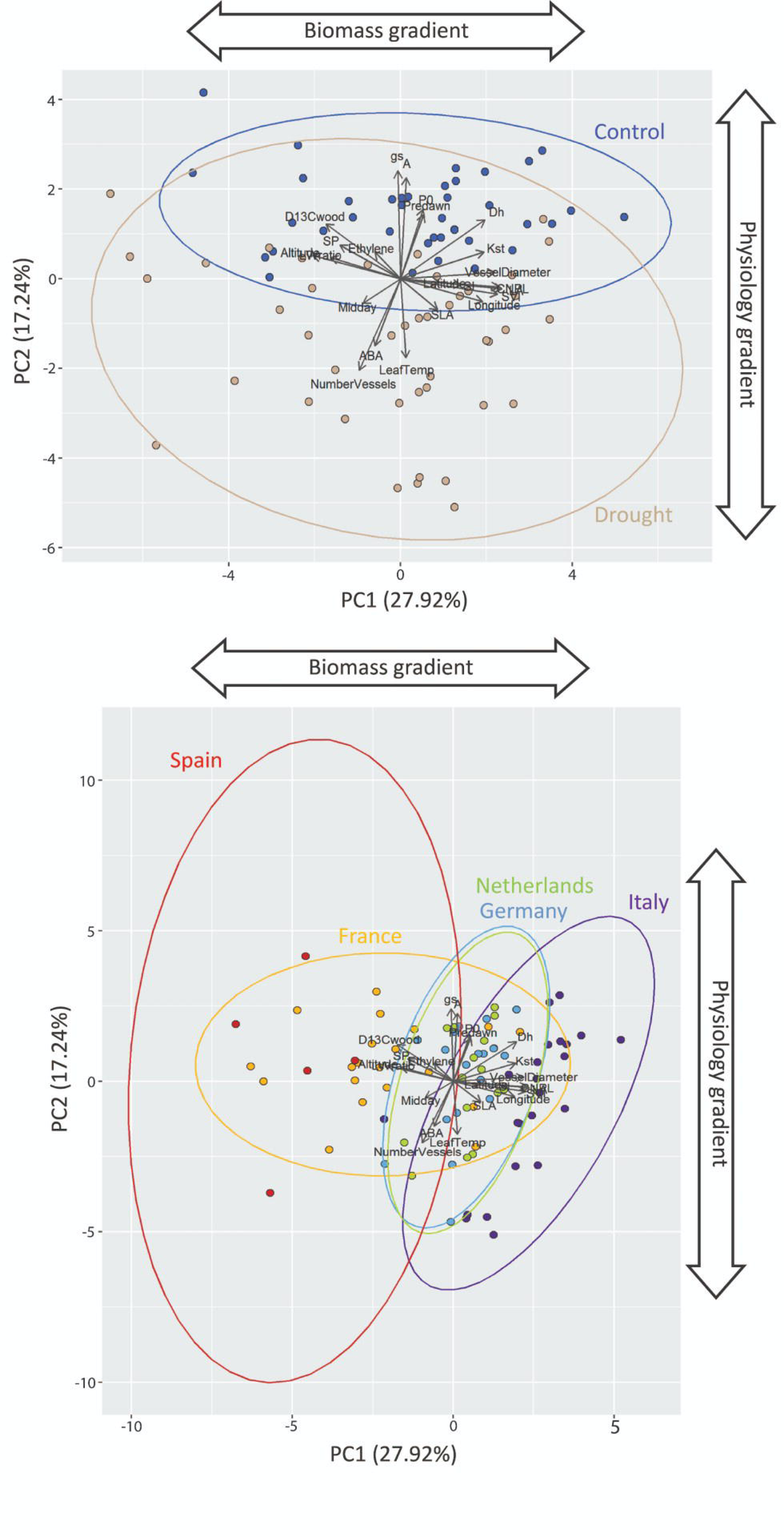
Principal Component Analysis of physiological and morphological traits. Grouped by irrigation regime (A) and genotype (B). g_s_, stomatal conductance (mmol m^-2^ s^-1^); A, CO_2_ assimilation (μmol m^-2^ s^-1^); P0, osmotic potential at full turgor (MPa); Predawn, predawn water potential (MPa); Midday, midday water potential (MPa); Dh, hydraulic weighted vessel diameter (µm); Kst, theoretical hydraulic conductivity (Kg s^-1^m^-1^ Mpa^-1^); VesselDiameter, wood vessel diameter; CNPL, cell number per leaf; SVI, stem volume index (cm^3^); SI, stomatal index (%); Latitude, latitude; Longitude, longitude; SLA, specific leaf area (cm^2^ g^-1^); Leaf Temp, leaf temperature (°C); NumberVessels, number of wood vessels (N mm^-2^); ABA, abscisic acid concentration (ng g^-1^ DW); SP, saccharification potential (% ODW); Ethylene, ethylene concentration (nl h^-1^ g^-1^); LWratio, leaf length:width ratio; D13C, wood carbon isotope discrimination (^0^/_00_); Altitude, altitude (m asl).

### Drought sensitivity

In general, drought decreased midday water potential and individual leaf area (r=-0.58, p<0.05, Figure 5A). While some genotypes such as Sp1, Ge4, Ne1 and It2 maintained, or increased, leaf growth under drought conditions, other genotypes, such as It4, Ne3, Fr2, It1 and Fr4, were identified as drought sensitive. Those genotypes with increased leaf size had less negative midday water potentials under drought. Simultaneously, reduced water availability was related to increased ABA levels, reduced stomatal index (r=0.56, p<0.05, Figure 5B) and more negative osmotic potential at full turgor (plotted as a percentage response of a negative value, r=0.50, p<0.05, Figure 5C). The genotypes which were identified as tolerant in terms of their individual leaf growth response, all showed intermediate levels of ABA compared to other genotypes with a maintenance or reduction in stomatal index and a low decrease in osmotic potential at full turgor. The exception to this was Sp1 which showed a high decrease in osmotic potential at full turgor in response to drought.

**Figure 5.**
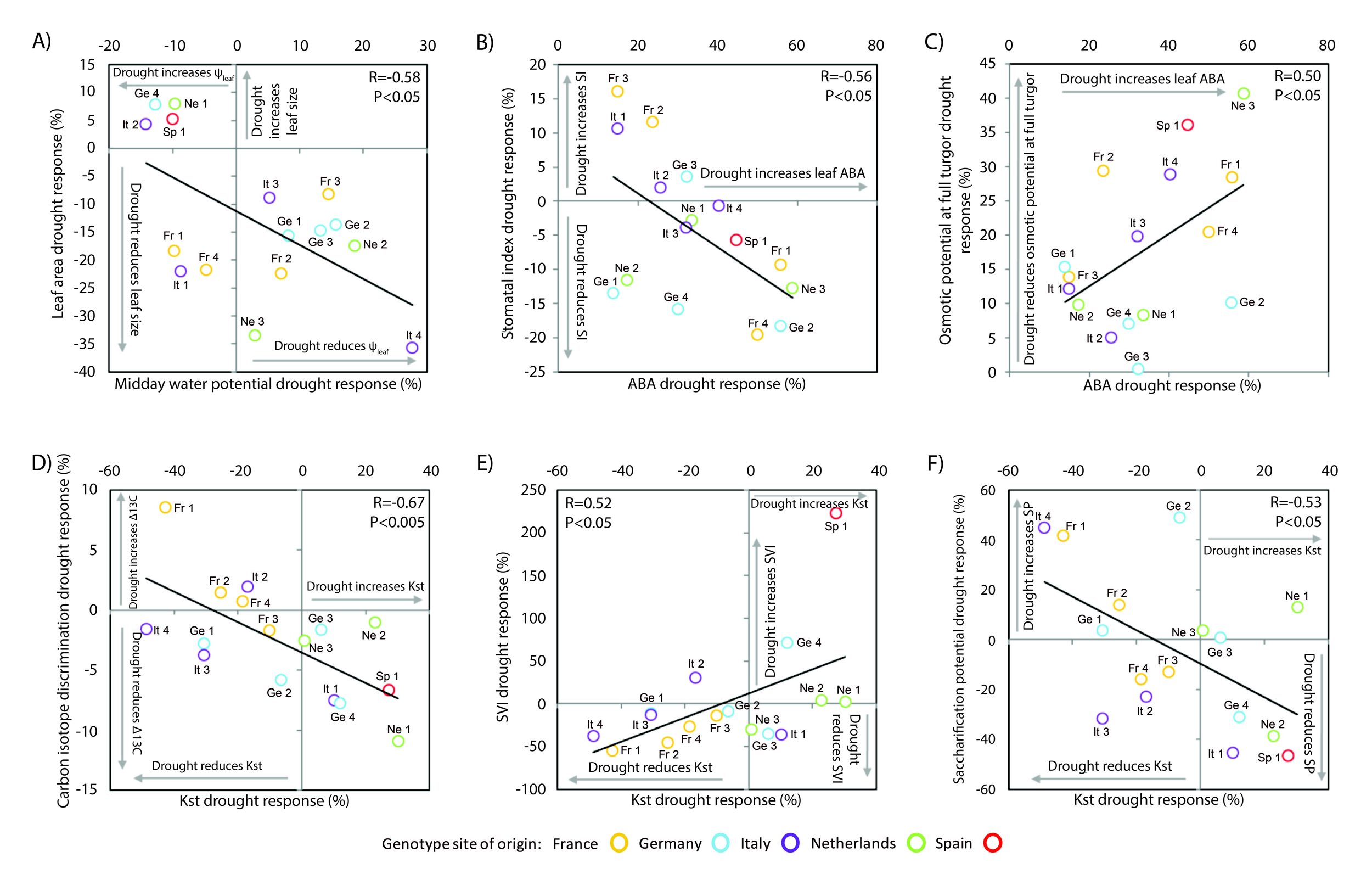
The drought response (percent difference between control and drought) of the diversity panel. Midday water potential versus leaf area (A), abscisic acid concentration versus stomatal index (B), abscisic acid concentration versus osmotic potential at full turgor (C), theoretical hydraulic conductivity versus wood carbon isotope discrimination (D), theoretical hydraulic conductivity versus stem volume index (E), theoretical hydraulic conductivity versus saccharification potential (F).

Highly productive trees were characterised by high K_st_ in both drought and irrigated regimes (Figure 3B–C). Further, in response to reduced water availability, increased K_st_ was associated with reduced Δ^13^C_wood_ (r=-0.67, p<0.005, Figure 5D), increased SVI (r=0.52, p<0.05, Figure 5E) and decreased saccharification potential (r=-0.53, p<0.05, Figure 5F). In general, the trees which maintained leaf growth under drought, also maintained or increased SVI. Further, these tolerant genotypes showed the most positive K_st_ responses under drought, linked to reduced Δ^13^C_wood_, and saccharification potential and improved SVI performance under drought.

To assess drought tolerance, the DRI and YSI of each genotype were compared (Figure 6). Eight genotypes had a YSI of SVI exceeding 1, indicating high yield under drought compared to the rest of the population (It3, It2, It1, Ge1, Fr3, It4, Ne2, Ne1, Figure 6A). Of these, five (It1, Fr3, It3, It4, It2) also showed higher K_st_ than the population when exposed to drought (Figure 6B). The Δ^13^C_wood_, and implied WUE of these genotypes, tended to be relatively sensitive to drought, with the exception of It1 and Ne1 (Figure 6C). Although, some of these genotypes were sensitive to drought when assessed via the DRI, they performed favourably when the YSI was considered. For example, It2 shows one of the largest reductions in Δ^13^C_wood_ (Figure 5D), yet, in absolute terms, it exhibits the lowest Δ^13^C_wood_ values (and highest WUE) of the population both under control and in response to drought. However, these low Δ^13^C_wood_ (high WUE), high yielding trees also demonstrate low saccharification potential (Figure 6D).

**Figure 6.**
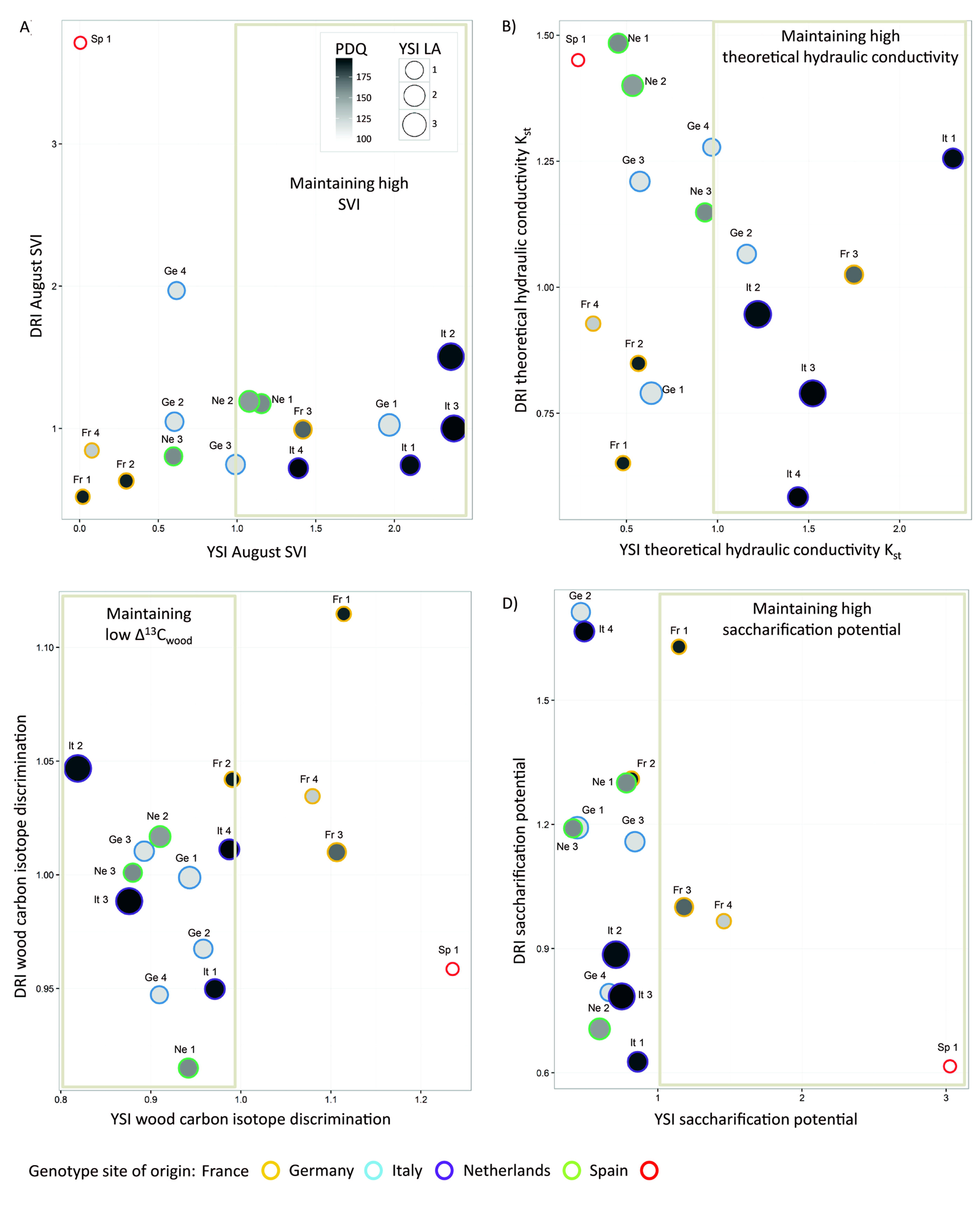
Drought resistance index (DRI) and yield stress index (YSI) comparisons. The size of the circular data point represents the yield stress index leaf area (YSI LA), with the grey scaling indicating the precipitation of the driest quarter (PDQ). Stem volume index (SVI) (A), theoretical hydraulic conductivity (B), wood carbon isotope discrimination (C) and saccharification potential (D).

## Discussion

This study identified significant natural genetic variation in a number of morphological traits in black poplar, as well as a high level of phenotypic plasticity across European sites and in response to drought. Further, this important model species exhibited wide variability in adaptive physiological drought responses and consequent productivity and saccharification potential. Taken together, these results have highlighted parallels and trade-offs between yield, WUE and saccharification potential as targets for breeding tolerant bioenergy trees for future climates.

The main source of variance between genotypes was morphological, with genotypes from areas of higher precipitation tending to be more productive, associated with larger, deltoid-shaped leaves comprising more individual cells. These genotypes (from Italy, Germany and The Netherlands) had a high theoretical xylem hydraulic efficiency with high stomatal index. Conversely, genotypes originating from drier regions (southern France and Spain) were less productive with smaller, rhomboid-shaped leaves consisting of fewer larger cells, lower stomatal index, and a reduced hydraulic efficiency (K_st_, D_h_). These differences in growth traits are considered to be adaptations to drought conditions arising from conditions at the site of origin (Regier *et al*. 2009; Cocozza *et al*. 2010; Viger *et al*., 2016). Although precipitation at the site of origin was not the main driver of leaf growth in this study, individual leaf size was tightly linked to longitude and altitude, implying a combination of meteorological factors influence adaptation. These morphological traits of genotypes from arid regions constrained water use, which might determine survival during episodes of intense and prolonged drought. Petiole xylem features are well correlated with leaf size in poplar, with large leafed genotypes containing larger vessels (Gebauer *et al*., 2016), which can transport more water (Brocious and Hacke, 2016). This coordination of vessel morphology and leaf anatomy may explain the relationship between genotype, biomass accumulation and climate of origin as in *P. tremula* and *P. tremuloides* (Hajek *et al*., 2014). Furthermore, the narrow vessels and small leaves of genotypes from drier regions may increase resistance to cavitation (Guet *et al*., 2015; Hajek *et al*., 2014; Hacke *et al*., 2010; Schreiber *et al*., 2015). Although these adaptations facilitate survival in water-limited environments, they also limit biomass accumulation in optimal and more moderate drought conditions. Thus, this adapted phenotype may be of limited value for high productivity in Europe if moderate droughts remain the status quo. Rather, our attention is drawn to the phenotypic plasticity displayed by the high yielding trees and under what conditions productivity ceases to be a useful selection trait.

Capturing this phenotypic plasticity is important and, in this study, leaf development seems critical. Leaf size was highly plastic in response to the growing environment in large leafed genotypes, such as those originating from northern Italy, which were responsive to drought when compared to small leafed genotypes (Viger *et al*., 2016). Individual leaf size is a well-established proxy of tree productivity (Rae *et al*. 2004, 2009; Monclus *et al*. 2005; Viger *et al*., 2016), as leaves are the main site for radiation interception, gas exchange and photosynthesis. While studies generally focus on a unique panel of genotypes grown in a single environment, here we have shown that the relationship between individual leaf area and yield is strongly independent of environment and is a highly robust trait for future selection.

In contrast, there was no consistent relationship between Δ^13^C and yield across sites, due to a genotype x environment interaction, similar to that reported for *P. deltoides* hybrids (Dillen *et al*., 2011; Chamaillard *et al*., 2011). In Italy, Δ^13^C and biomass growth were negatively correlated as in previous studies with this species (Viger *et al*., 2016; Guet *et al*., 2015) and *P. trichocarpa* (Guerra *et al*., 2016), while in Belgium the relationship was absent as in *P. deltoides* x *P. nigra* hybrids (Monclus *et al*., 2005; Marron *et al*. 2005, Fichot *et al*., 2010). Therefore, genotypes with high productivity showed the highest WUE in Italy, and maximised the difference in WUE between sites. This is consistent with previous results in black poplar (Viger *et al*., 2016) and other tree species (Zhang *et al*., 1997) and is possibly due to drought avoidance strategies in genotypes adapted to dry environments. While genotypes from arid regions are adapted to low precipitation during the dry season, they may capitalise on periods of water availability such as during these irrigated experiments through maximising carbon gain with associated water losses. This idea is supported by the high stomatal conductance levels measured in the Spanish genotype at the Italian field site, coupled with low values of WUE. Genotypes from wetter climates with high productivity adjusted their physiology to maintain high rates of photosynthesis in warmer climates, without increasing stomatal conductance to the same degree, thereby increasing WUE. The lower plasticity in WUE of genotypes from drier climates aligns with a conservative use of resources of plants adapted to stressful environments (Valladares *et al*., 2007).

Genotypic variability of physiological traits was much lower than for morphological growth traits, indicating stronger environmental control and lower heritability of physiology (Rae *et al*., 2004, 2009). In general, physiological and biomass traits were mostly decoupled, as previously reported in *Populus* (Monclus *et al*., 2005; Fichot *et al*., 2010) where instantaneous photosynthesis rate is not always directly related to biomass production (Soolanayakanahally *et al*., 2009). The small variability in physiological traits compared to the large genotypic differences in morphological traits, suggests that a key driver of tree productivity is xylem hydraulic efficiency, which is, in turn, attributable to water availability at the site of origin, and enables the development of many large leaves to drive biomass accumulation (Marron *et al*., 2006). In support of this hypothesis, high water transport capacity has been correlated with biomass accumulation in both *Populus* and *Salix* species (Fichot *et al., 2009;* Hajek *et al*., 2014; Gebauer *et al*., 2016). The link between this increased water transport capacity and the increased production of cells and larger leaves is key to devising strategies for selection and breeding for future European environments.

However, high productivity was at the expense of saccharification potential under drought conditions. Previously, glucose release has been shown to be positively related to wood density (Serapiglia *et al*., 2013), and both negatively related (Acker *et al*., 2014), or unrelated, to lignin content (Wildhagen *et al*., 2017). Yet in this panel of diverse genotypes originating from across Europe, saccharification potential does not seem related to wood density. This trade-off between yield and saccharification potential may be minimal given yield variability is far higher than that for saccharification potential, with moderate saccharification potential combined with high productivity being favoured when compared to low yielding, high saccharification potential trees. Further, genotypes such as It2 were high yielding and, despite showing one of the largest reductions in Δ^13^C_wood_ in response to drought, exhibited the highest WUE of the population under control, and in response to drought. This dual-target of yield and WUE is likely more important than selecting for high saccharification potential alone.

Underlying these wood traits and yield effects, trees adopt variable drought response strategies. In general, drought reduced stomatal conductance and leaf size, consistent with higher leaf ABA concentration and lower midday water potential. Further, genotypes which accumulated more ABA, achieved lower SIs than low ABA trees. While midday water potential declined in response to drought in most genotypes, with variable leaf growth impacts, some genotypes maintained or improved leaf growth under drought by increasing midday water potential. These drought tolerant genotypes had intermediate ABA levels compared to other genotypes, with decreased osmotic potential at full turgor. Although drought increased leaf growth of Sp1 (alongside a moderate increase in ABA), it showed a much larger reduction in osmotic potential at full turgor than other tolerant genotypes. Nevertheless, drought increased ABA concentrations and lowered osmotic potentials at full turgor, but these changes were not consistently related to yield. Genotypic variation in osmotic adjustment was not linked to the climate of origin and our findings suggest that using osmotic potential at full turgor to predict genotypic variation in drought tolerance within a species might not be useful in *Populus*.

Further, these putative drought tolerant genotypes maintained or reduced SI, while others reduced SI alongside significant leaf growth defects. Moderation of stomatal patterning is an alternative mechanism by which plants can regulate water loss in response to environmental change (Casson and Hetherington, 2010; Franks *et al*., 2015; Silva *et al*., 2009). The relationship found here suggests different genotype-specific strategies for stomatal control in *P. nigra*, either by reducing the number of stomata or increasing ABA concentration. These two strategies might have converged into conserved isohydric behaviour. This indicates a diversity of mechanisms allowing tree adaptation to different environments, which may consequently exhibit different sensitivities to ABA (Negin and Moshelion, 2016), or hydraulic adjustment strategies, to maintain cell turgor. Genotypes originating from drier environments regulated their stomata more loosely, while high yielding genotypes from wetter climates demonstrated high stomatal control and drastically reduced stomatal conductance under drought despite higher predawn water potentials. This may be supported by larger and deeper root systems (Monclus *et al*., 2005; Viger *et al*., 2016). At the same time, low values of stomatal conductance were coupled with higher WUE, low vessel density and high D_h_, reinforcing the idea that highly productive genotypes are drought avoiders and respond rapidly to drought with larger root systems and enhanced capacity to reduce water loss.

Using the natural genetic diversity of *P. nigra*, we have elucidated adaptive and plastic drivers of productivity, wood quality and drought tolerance in this fast-growing temperate tree species. Further, we have highlighted the physiological and morphological drivers which underlie these plastic and adaptive responses to drought as important traits for both breeders and growers in a future, water-scarce environment. Highly productive genotypes, originating from less arid regions, combined increased vigor with extensive phenotypic plasticity, when grown across three contrasting European environments and showed high YSI values in response to drought. Vigor was characterized by high theoretical xylem hydraulic conductivity and WUE, rapid leaf cell production and large leaves. These vigorous, high productivity genotypes seem best suited to multiple droughted- and well-watered European environments. Moreover, although breeding for WUE and saccharification potential may be complicated by higher environmental control of these traits, we have identified candidate genotypes which combine these desired traits for future breeding of bioenergy trees to ensure resilience for future climates.

## Supplementary data

Figure S1. Rainfall and temperature were monitored at the northern Italian field site, with the grey area indicating the period when the drought was applied (A). Using a Before and After Control Index (B.A.C.I.) design, the drought treatment was shown to significantly reduce soil moisture content at both 20 and 40 cm depths (* for p value<0.05, ** for p value<0.01) (B).

Table S1. All physiological and morphological traits show distinct correlations or independence when the diversity panel was grown in northern Italy: control and drought pooled (A), control (B), and drought (C).

## Acknowledgements

This work was conducted in the framework of the project WATBIO (Development of improved perennial biomass crops for water-stressed environments), which is a collaborative research project funded from the European Union’s Seventh Programme for research, technological development and demonstration under grant agreement No. 311929. J H B-B acknowledges funding from the Amar-Franses and Foster-Jenkins Trust through the award of a scholarship.

We thank Simon Rüger and Aneta Bargiel (YARA ZIM Plant Technology GmbH, Hennigsdorf, Germany) for their collaboration. We acknowledge the providers of the original *P. nigra* genotypes, which were collected in the European Commission project EVOLTREE (contract No. 016322, Directorate General Research within the Sixth Framework for Research as part of the Network of Excellence) and Catherine Bastien (INRA, Orléans, France) for supplying the stock cuttings.

## Notes

### Competing Interest Statement

The authors have declared no competing interest.

