## Supplementary figures and images for "The ideotype for drought tolerance in bioenergy *Populus nigra*"

### Supplemental Figure 1

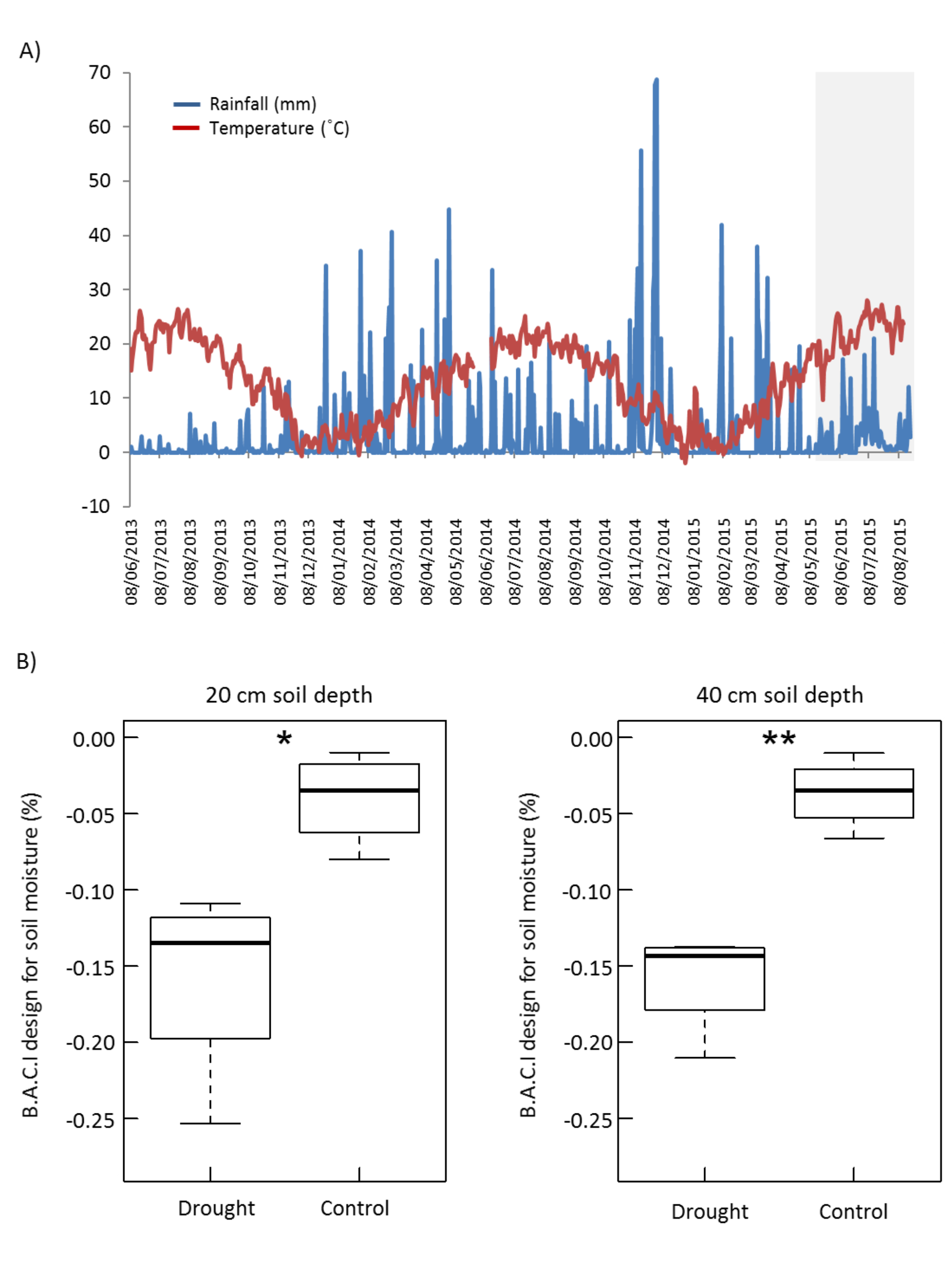
